# Antagonistic center-surround mechanisms for direction selectivity in the retina

**DOI:** 10.1101/831453

**Authors:** Lea Ankri, Elishai Ezra-Tsur, Shir R. Maimon, Nathali Kaushansky, Michal Rivlin-Etzion

## Abstract

A key feature in sensory processing is center-surround receptive field antagonism. Retinal direction-selectivity (DS) relies on asymmetric inhibition from starburst amacrine cells (SAC) to direction selective ganglion cells (DSGC). SAC exhibit antagonistic center-surround, depolarizing to light increments and decrements in their center and surround, respectively, but the role of this property in DS remains elusive. We found that a repetitive stimulation exhausts SAC center and enhances its surround and used it to distinguish center-from surround-mediated responses. Center, but not surround stimulation, induced direction-selective responses in SAC, as predicted by an elementary spatiotemporal model. Nevertheless, both SAC center and surround elicited direction-selective responses in DSGCs, but to opposite directions. Physiological and morphology-based modeling data show that the opposed responses resulted from inverted DSGC’s excitatory-inhibitory temporal balance, indicating that SAC response time rules DS. Our findings reveal antagonistic center-surround mechanisms for DS, and demonstrate how context-dependent center-surround reorganization enables flexible computations.

## Introduction

Antagonistic center-surround receptive field organization is a common motif in the computation of sensory inputs as it enables higher discrimination of sensory information (Bastian et al., 2002; Hubel and Wiesel, 1962; Knudsen and Konishi, 1978; Yokoi et al., 1995). In the visual system, On-center cells depolarize to light increments in the center of their receptive fields, but are inhibited by light increments in their surround. Off-center cells display a similar center-surround antagonism, with opposite polarity preference. These cells frequently exhibit surround activation: a depolarizing response to a stimulus of the non-preferred polarity in the periphery (Barlow, 1953; Kuffler, 1953). Antagonistic center-surround receptive fields are present at all the levels of visual processing, starting from the retina up to the thalamus and visual cortex and are considered a key mechanism for the enhancement of edge detection and color vision (Dacey, 1999; Marr and Hilderth, 1980; Usrey and Alitto, 2015). This spatial antagonism depends on the nature of the stimulus and has the property of dropping out in dim illumination and strengthen at high light levels (Barlow et al., 1957; Bisti et al., 1977; Dedek et al., 2008; Farrow et al., 2013; Rivlin-Etzion et al., 2018). Thus, center-surround organization is not fixed, qualitatively changing the cell’s function under distinct sensory contexts.

In the retina, On-Off direction selective ganglion cells (DSGC) respond maximally to motion in their preferred direction (PD) and minimally to motion in the opposite, null direction (ND). Starburst amacrine cells (SAC) mediate the directional response via asymmetric wiring, as only processes oriented in the DSGC’s null direction tend to form inhibitory synapses on its dendrites (Figure 1A) (Briggman et al., 2011). Although SAC somatic voltage shows no directional preference to linear motion, its processes act as autonomous units with directional preference, responding with greater depolarization to motion away from the cell soma (centrifugal; CF) than to motion towards cell soma (centripetal; CP) (Ding et al., 2016; Euler et al., 2002; Fransen and Borghuis, 2017; Koren et al., 2017). Due to their wiring specificity, SAC processes that innervate a given DSGC display directional preference to its null direction, which is expected to result in stronger GABA release during null motion than during preferred motion. This asymmetric inhibition is thought to play a crucial role in direction selectivity.

**Figure 1.**
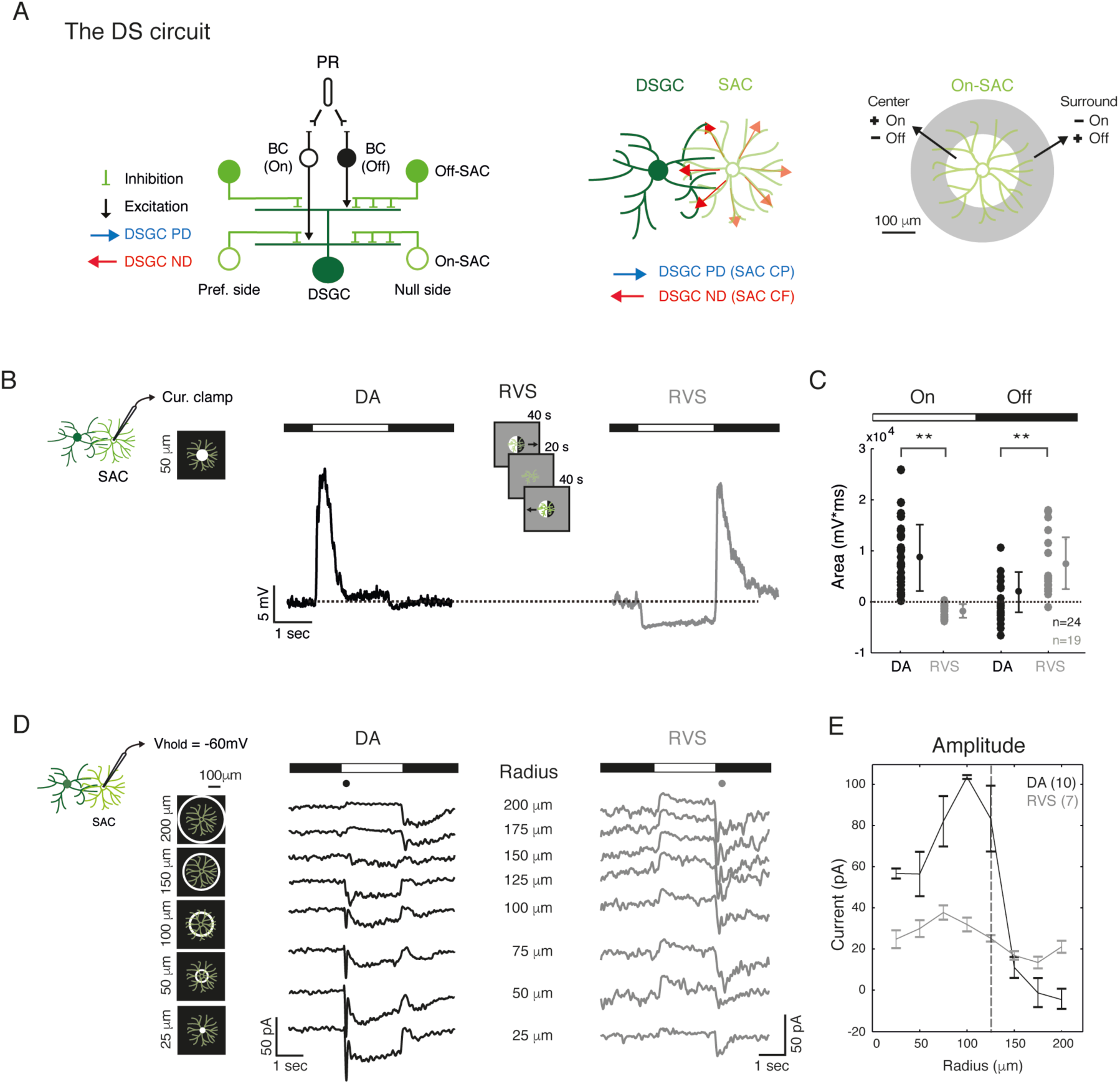
RVS reorganizes SAC center-surround receptive field. *(A) Left*: Schematics of the direction selective (DS) circuit, side view. SAC on the null side form more inhibitory connections with the DSGC than SAC on the preferred side. Excitation from On- and Off-BC to On- and Off-SAC is not shown for simplicity. *Middle*: Schematics of a DSGC with one innervating SAC, top view. DSGC’s null motion corresponds with CF motion of its innervating SAC processes, and DSGC’s preferred motion corresponds with their CP motion. *Right*: schematics of On-SAC antagonistic center-surround organization. PR – photoreceptors; BC - bipolar cell; PD – preferred direction; ND – null direction; CF – centrifugal; CP – centripetal. (B) Average current-clamp recording of an example On-SAC in response to a 2 sec static bright spot (illustrated on the left, 50 μm radius; 10 repetitions) in dark adapted conditions (DA, black) and following RVS (RVS, grey). RVS is illustrated in center. (C) Area under the curve for 1.5 sec of On-SAC responses to light onset and to light offset in dark adapted conditions and following RVS. Group means and STD are indicated on the side by circles and error bars; Asterisks indicate statistical significance (**p < 0.001). n denotes number of cells. Dashed line in (B) and (C) denote the baseline voltage and area, respectively. (D) Excitatory current recordings (V_hold_ = −60 mV; averaged over 4 repetitions) from two example On-SAC in response to 2 sec static rings of 8 different radii (5 of which are illustrated on left), in dark adapted conditions (DA, black) and following RVS (RVS, grey). RVS is as shown in (B). The rings’ radii are specified in the center next to the corresponding traces. Black and grey dots denote the period used for the population analysis in (E). (E) Maximal amplitude (mean ± STD) of excitatory current as a function of ring radius in dark adapted conditions (acquired during ring onset; black) and following RVS (acquired during ring offset; grey). Dashed vertical line denotes the distal boundary of SAC excitatory receptive field (Ding et al., 2016; Vlasits et al., 2016). Numbers of cells in each group are in brackets.

Neurons in the direction selective circuit display center-surround receptive field organization, but if and how this organization contributes to the computation of motion direction remains controversial. SAC were shown to have excitatory inputs confined to their proximal 2/3 dendritic arbors, and it was hypothesized that this center organization contributes to CF preference (Ding et al., 2016; Vlasits et al., 2016). Reciprocal SAC-SAC lateral inhibition enhances CF preference (Lee and Zhou, 2006; Zhou and Lee, 2008), but this surround inhibition is not strictly required as blocking GABA receptors fails to fully eliminate SAC directional responses (Hanson et al., 2019; Hausselt et al., 2007; Oesch and Taylor, 2010). While the role of surround inhibition in direction selectivity was often tested by abolishing surround responses, it was never tested by abolishing center responses.

We previously showed that a short repetitive visual stimulation (RVS) switches polarity preference in SAC (Vlasits et al., 2014). Here, we find that this occurs due to reorganization of SAC receptive field by RVS, which abolishes center and enhances surround activation. We then employ RVS-evoked changes to decipher the role of SAC center and surround organization in direction selectivity. RVS abolished SAC CF preference. Yet, both center-mediated activation (assessed prior to RVS) and surround-mediated activation (assessed following RVS) elicited directional responses in DSGC: center activation elicited responses to the preferred direction, while surround activation elicited response to the null direction. These opposed direction responses were mediated by SAC response timing. Thus, we demonstrate that SAC center-surround organization rules DSGC computation by controlling the timing of inhibition rather than its amount. Thereby, we reveal a novel concept for center-surround antagonism in the retinal direction selective circuit, where PD motion is encoded in the center receptive field and ND motion in the surround.

## Results

### On-SAC lose center and enhance surround activation following RVS

To dissect the network mechanisms underlying direction selectivity in the retina, we characterized the response properties of SAC using targeted recordings from somata of On-SAC which are fluorescently labeled in mGluR-EGFP mice (Watanabe et al., 1998). We previously showed that a short repetitive visual stimulation (RVS) with drifting gratings in the time scale of minutes can switch polarity of excitatory input to SAC in response to a static bright spot stimulus (Vlasits et al., 2014). Here, we confirmed these results using current-clamp recordings, demonstrating that On-SAC depolarize to the onset of a bright spot in dark-adapted conditions, but depolarize to the spot disappearance following RVS (Figure 1B, C). We then sought to examine the origin of polarity switch in On-SAC. Since opposite polarity preferences emerge in center and surround receptive fields of many retinal neurons, we set to search for possible changes in SAC receptive field organization following RVS.

To characterize On-SAC excitatory receptive field, we recorded their excitatory currents (V_hold_ = −60 mV) while presenting them with eight static rings of different radii (25-200 μm; Figure 1D). Excitatory inputs to dark adapted On-SAC increased with ring radius, reaching a maximum value in response to ∼100 μm radius. For radii bigger than 100 μm the excitatory currents amplitude rapidly decreased, showing no excitation in response to rings larger than 175 μm radius (Figure 1D, E). These findings match the distribution of glutamatergic inputs from bipolar cells to SAC, which are restricted to the inner 2/3 of the dendritic tree (Ding et al., 2016; Vlasits et al., 2016). Surround activation was observed in a portion of the cells, where rings of radii larger than 150 μm revealed loss of excitation to ring onset accompanied by increased excitation upon ring offset (Figure 1D) (Lee and Zhou, 2006). Following RVS, excitation to On-SAC emerged in response to the rings offset in all ring’s radii (Figure 1D), reflecting the polarity switch observed in SAC response to central spot activation (Figure 1B). These Off inputs to On-SAC showed little dependency on ring radius, indicating a loss of the characteristic proximal excitation to light onset and center-surround organization (Figure 1E). The change in On-SAC excitatory receptive field following RVS probably results from the activation of surround circuits in the outer retina (see *Discussion*). Since SAC proximal excitation is thought to contribute to their CF preference (Ding et al., 2016; Vlasits et al., 2016), loss of proximal excitation may affect SAC directional preference. To explore this possibility, we investigated SAC directional responses in dark-adapted conditions and following RVS.

### On-SAC lose their directional preference and their responses are shifted in time following RVS

We presented On-SAC with expanding (CF motion) and collapsing (CP motion) drifting rings centered on the cell’s soma while recording SAC somatic voltage in current-clamp mode (Figure 2A). In accordance with previous reports, dark adapted (DA) On-SAC displayed stronger depolarization in response to CF than to CP motion (Figure 2A, B, top) (Ding et al., 2016; Euler et al., 2002; Fransen and Borghuis, 2017; Koren et al., 2017). The CF preference was reflected by a positive annular direction selective index (A-DSI; see *Methods*; A-DSI_DA_ = 0.19 ± 0.2; Figure 2C). Responses to CF and CP motion did not only differ in amplitude but also in kinetics, displaying shorter rise time in response to CF motion, as quantified by the latency from response onset (set to the time of 20% the peak value) to the time of response peak (rise_time_DA__CF = 75 ± 38 ms, rise_time_DA__CP = 162 ± 71 ms; p < 0.001; Figure 2D; for additional measures see Supplementary Figure 4).

**Figure 2.**
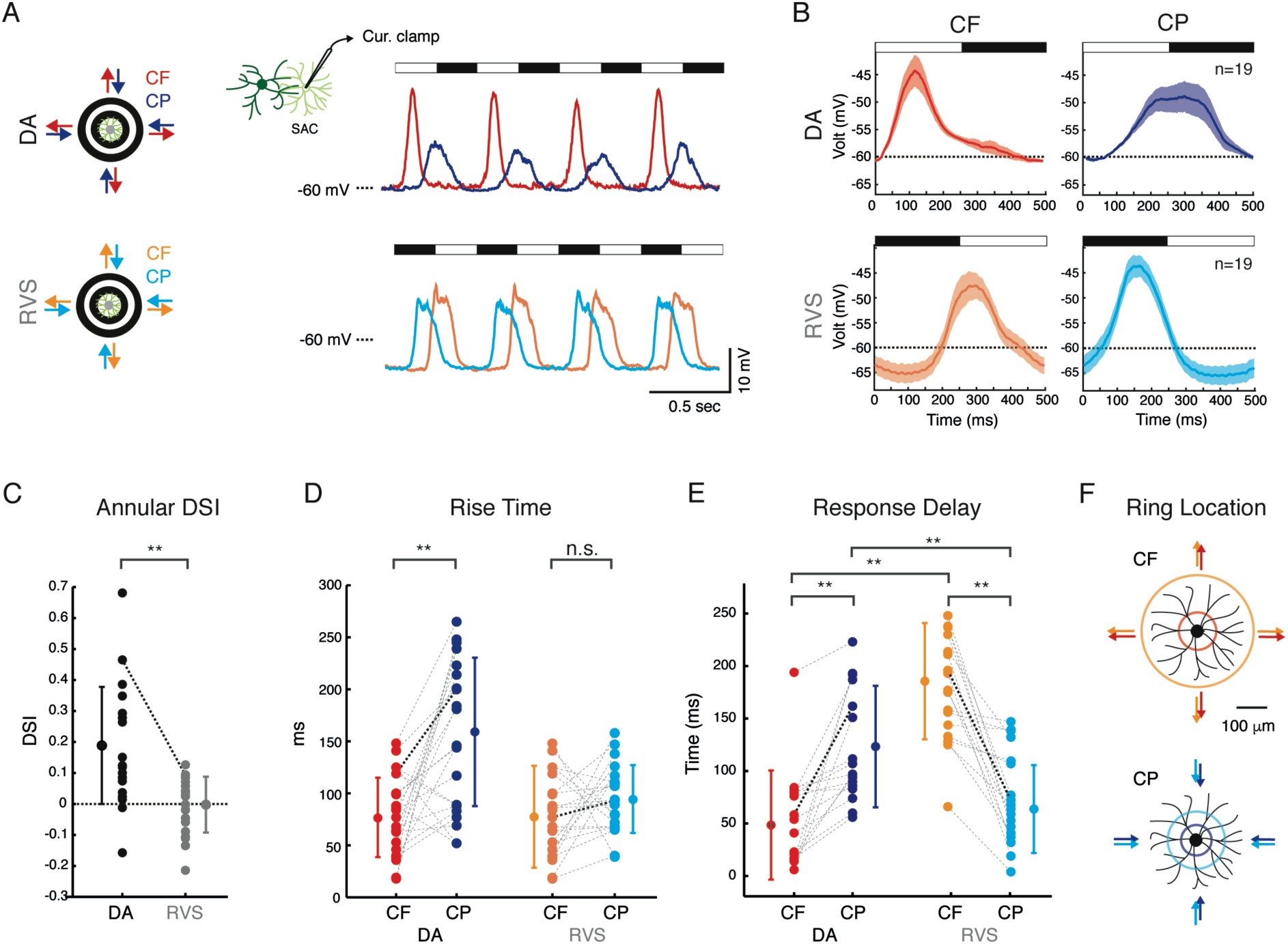
RVS abolishes SAC CF preference and shifts its response time. *(A) Left*: illustration of drifting rings stimulus. *Right*: Average current clamp recordings from an example On-SAC in response to CF and CP motion (5 repetitions) in dark adapted conditions (DA, top) and following RVS (RVS, bottom). Black and white horizontal bars denote the timing of dark and bright rings on the cells’ processes with respect to the soma. (B) One cycle waveforms of On-SAC population responses to CF and CP rings (average ± SEM) in dark adapted conditions (top) and following RVS (bottom). Horizontal dashed line in (A) and (B) denote −60 mV voltage. (C) Annular DSI of On-SAC population, calculated based on response amplitudes for CF vs. CP motion in dark adapted conditions and following RVS. (D) Rise time of CF and CP motion responses, measured as the latency from response onset to response peak in dark adapted conditions and following RVS. (E) Response delay during CF and CP motion, measured as the latency from the time the bright and dark rings appeared on the cell’s receptive field to response onset in dark adapted conditions and following RVS, respectively. (F) Colored rings depict the location of the ring’s leading edge relative to SAC (white ring for dark adapted, black for RVS) at the time of average response onset depicted in (E). For (C-E), group means and STD are indicated on the side by circles and error bars; dashed lines connect between values of the same cell; bold dashed lines represent the example cell shown in (A). Asterisks indicate statistical significance (**p < 0.001).

Following RVS, On-SAC response asymmetries disappeared, revealing similar voltage waveforms during CF and CP motion (Figure 2A, B, bottom). This was reflected in reduced A-DSI values which clustered around zero (A-DSI_RVS_= 0.01 ± 0.09; A-DSI_DA_ vs. A-DSI_RVS_: p < 0.001; Figure 2C) and similar response kinetics (rise_time_RVS__CF = 77 ± 34 ms, rise_time_RVS__CP = 95 ± 29 ms; p = 0.19; Figure 2D; Supplementary Figure 4). On-SAC enhanced hyperpolarization that emerged in response to a stationary light onset following RVS (Figure 1B) was also detected in response to the drifting rings (Figure 2A, B).

A careful examination of SAC response waveforms demonstrated that they shifted in time following RVS, with responses to CF and CP motion shifting in opposite manners. To quantify this, we measured the time of response delay as the time from stimulus onset to response onset. For dark adapted SAC, we assessed the stimulus onset as the time the leading bright edge of the ring encountered the cell’s dendritic arbor (i.e., when leading edge traversed most proximal dendrites for CF motion and most distal dendrites for CP motion). Following RVS, we used similar measures but in relation to the dark edge of the ring, due to SAC polarity switch. We found that response delay to CF motion increased on average by 128 ms following RVS (response_delay_DA__CF = 49 ± 52 ms, response_delay_RVS__CF = 177 ± 53 ms; p < 0.001; Figure 2E). The mean response timings corresponded to the leading edge location at 43 and 159 μm away from cell soma in dark adapted condition and following RVS, respectively (Figure 2F, top). For CP motion, response delay was shortened on average by 59 ms following RVS (response_delay_DA__CP = 123 ± 58 ms, response_delay_RVS__CP = 64 ± 42 ms; p < 0.005; Figure 2E). The mean response timings corresponded to the leading edge location at 110 and 57 μm away from the cell’s distal processes in dark adapted condition and following RVS, respectively (i.e., 40 and 93 μm from cell soma; Figure 2F, bottom). These spatial measurements are consistent with the observed expansion of SAC excitatory receptive field (Figure 1D, E). SAC phase shift was also evident when assessed based on response peak time or on the argument of the Fourier transform (Supplementary Figure 1). Blocking GABA-A receptors with gabazine had no effect on response timing, although it abolished the enhanced hyperpolarization observed following RVS (Supplementary Figure 2A). Moreover, excitatory currents to SAC soma resembled SAC voltage responses and displayed a similar phase shift in response to drifting rings stimulation (Supplementary Figure 2B, C). These results suggest that excitation, rather than inhibition, underlies the observed shift in time of SAC responses following RVS.

### Center-surround reorganization explains phase shift in SAC responses and loss of CF preference

We hypothesized that SAC loss of center and enhancement of surround activation can underlie the phase shift in their responses to drifting rings. To test this possibility, we generated a morphology-based modelling environment and simulated SAC responses to drifting rings stimuli (see *Methods*). The morphology of the simulated SAC replicated that of an On-SAC which we filled with fluorescence dye and reconstructed (Figure 3A). The excitatory receptive field of the simulated SAC varied to mimic the experimental data. In the dark-adapted SAC simulation, excitatory receptive field was restricted to the proximal 2/3 of the dendritic arbor (Ding et al., 2016; Vlasits et al., 2016) whereas for the RVS simulation the excitatory receptive field was restricted to the distal 2/3 of the dendritic arbor (Figure 3A). In response to CF motion, the spatial shift of the excitatory receptive field caused a delay of ∼80 ms in depolarization of the post-RVS simulated SAC relative to the dark-adapted simulated SAC (Figure 3B). In response to CP motion, depolarization of the post-RVS simulated SAC preceded that of the dark-adapted simulated SAC by ∼40 ms (Figure 3B). These trends are in line with the phase shifts observed in our experimental data following RVS and could potentially shift the timing of SAC signaling to DSGC. Yet, SAC release sites are restricted to distal processes (Ding et al., 2016) and SAC somatic voltage might deviate from their dendritic voltage due to non-linearities in SAC dendrites (Hausselt et al., 2007; Lee et al., 2010; Oesch and Taylor, 2010; Ozaita et al., 2004; Xu et al., 2003). We therefore extracted the voltage in the distal processes of dark adapted and RVS simulated SAC and found a phase shift that is comparable to the one observed in the somatic voltage (Figure 3C). Hence, spatial organization of SAC excitatory receptive field predominantly controls the time of its directional response in a similar manner for the soma and for the distal dendrites.

**Figure 3.**
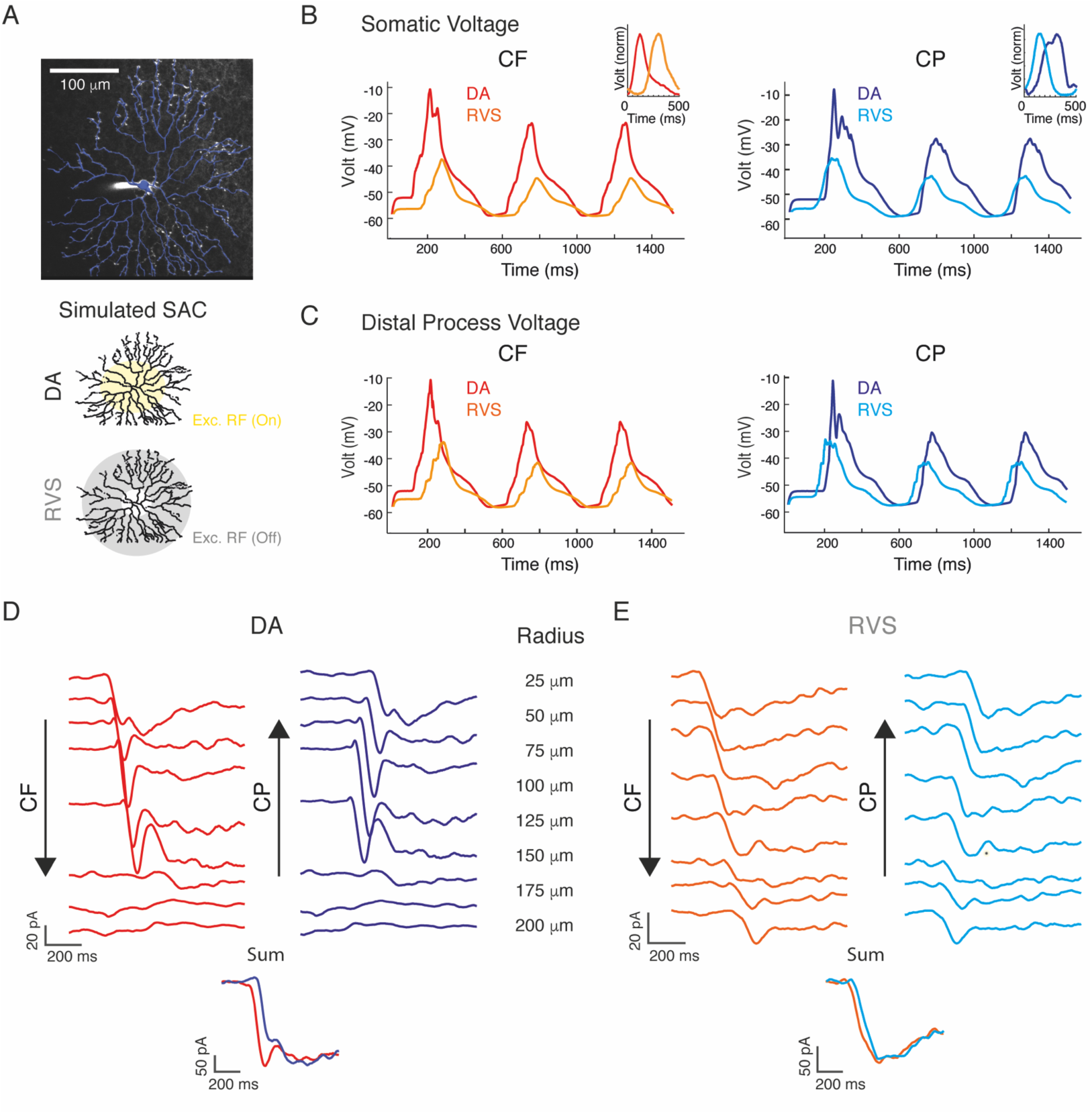
Excitatory center-surround receptive field organization dictates both SAC response time and directional preference. (A) *Top*: Projection of an On-SAC filled with fluorescent dye (white) and its reconstruction (blue) which was used for the morphology-based simulation. *Bottom*: Schematics of excitatory receptive field, covering 2/3 of the proximal dendrites in dark adapted conditions (top, yellow area) and 2/3 of the distal dendrites following RVS (bottom, grey area). (B) Example somatic voltage of a simulated SAC responses to CF and CP motion under the two excitatory receptive fields shown in (A). Dark adapted traces (red; blue) are plotted on top of RVS traces (orange; cyan). Insets depict the superimposed normalized average responses to CF and CP motion recorded from On-SAC. (C) Example dendritic voltage of a simulated SAC responses (d=135 μm from cell soma) to CF and CP motion in dark adapted conditions and following RVS. (D) Population average EPSCs (Vhold = −60 mV) evoked in dark adapted SAC in response to rings of different radii (noted on right). The traces are shifted in time to simulate soma-to-dendrite CF motion (left) and dendrite-to-soma CP motion (right). *Bottom*: Linear summation of the shifted responses. (E) As in (D), but for SAC recorded following RVS.

Next, we tested whether the observed symmetry in SAC response amplitude and kinetics following RVS can also be predicted from the change in their receptive field organization. We used SAC EPSCs evoked in response to static rings (Figure 1D, E) to simulate SAC response to motion by shifting responses in time in an orderly manner and summing them (Lien and Scanziani, 2018), either from smallest ring to largest one – to mimic CF motion, or from largest ring to smallest one – to mimic CP motion (Supplementary Figure 3). We set the delay to fit the velocity of the moving rings. We found asymmetries in the summation of EPSCs in dark adapted SAC, revealing faster response kinetics with larger amplitude for CF-like motion compared with CP-like motion (Figure 3D). Following RVS, the asymmetries in the response summation disappeared, revealing similar waveforms for CF- and CP-like motion (Figure 3E). This loss of asymmetry can be predicted from SAC homogenous responses to all rings’ radii following RVS (Figure 1D, E). Hence, a simple spatiotemporal model of the excitatory inputs to SAC can explain their responses both in dark adapted conditions and following RVS, emphasizing the tight control of SAC receptive field organization on their process’s directional preference.

### Loss of directional preference in SAC is reflected in the inhibitory inputs to DSGC

Thus far we showed that SAC receptive field organization can dictate time and directional preference of their motion responses. Thus, RVS provides us with a unique opportunity to decipher the role of SAC receptive field organization in the computation of motion direction in DSGC. One may expect that the loss of SAC directional responses would abolish direction selective responses in DSGC, but we previously showed that DSGC can reverse their directional preference following RVS (Rivlin-Etzion et al., 2012). To draw a possible link between the observed changes in SAC responses and reversal of directional preference in DSGC, we conducted cell-attached targeted recordings from posterior-preferring On-Off DSGC which are fluorescently labeled in Drd4-EGFP and TRHR-EGFP mice (Huberman et al., 2009; Rivlin-Etzion et al., 2011). We recorded the spiking activity and confirmed reversal of directional preference by measuring DSGC directional tuning in response to 3 sec of linear gratings drifting in 12 different directions in dark-adapted conditions and following RVS (Figure 4A).

**Figure 4.**
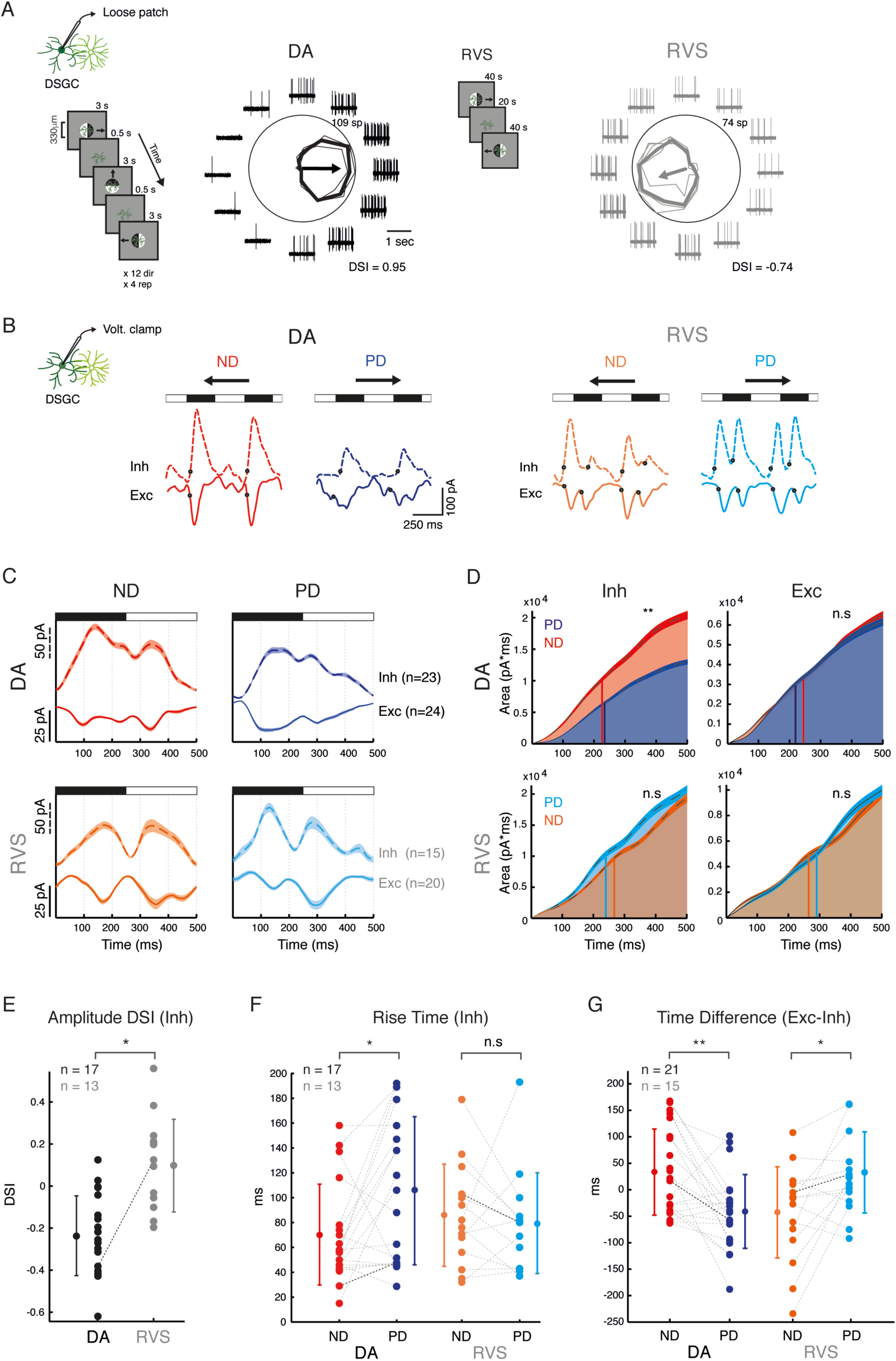
Time shift of inhibition relative to excitation is sufficient to reverse DSGC directional preference following RVS. (A) An example DSGC directional tuning to drifting gratings in 12 different directions (shown on the left) in dark adapted conditions (DA, black) and following RVS (RVS, grey). Traces are example of 1 sec recordings in each direction. Polar plots represent the mean response (spike count during 3 s) and thin lines show the responses in each repetition. Arrows represent the preferred direction and their length correspond to the normalized vectorial summation (with outer ring equal to 1). RVS is shown in the center. (B) Voltage clamp recordings of an example DSGC in response to gratings moving in the PD (blue, cyan) and ND (red, orange) in dark adapted conditions (left) and following RVS (right). Waveforms are averages of two cycles over 4 repetitions of inhibitory (dashed line; V_hold_ = 0 mV) and excitatory (continuous line; V_hold_ = −60 mV) currents; dark dots represent response onset, calculated as time of 20% of the maximal phase value. (C) One cycle average waveform ± SEM of the currents recorded from DSGC population during ND (left) and PD (right) motion, in dark adapted conditions (top) and following RVS (bottom). Numbers of cells are denoted in parentheses. For A&B, black and white horizontal bars denote appearance of dark and bright gratings relative to the cells’ soma. (D) Cumulative current during PD and ND motion of inhibitory (left) and excitatory (right) average waveforms plotted in (C). Vertical lines denote the time of 50% maximal value (PD: blue and cyan; ND: red and orange, for dark adapted conditions and following RVS, respectively). (E) DSI for inhibition based on current amplitude in dark adapted conditions and following RVS. (F) Rise time of inhibitory current during PD and ND motion, measured as the latency from response onset to response peak in dark adapted conditions and following RVS. (G) Time difference between excitatory and inhibitory response onset during PD and ND motion in dark adapted conditions (left) and following RVS (right). For (E-G), group means and STD are indicated on the side by circles and error bars; dashed lines connect between values of the same cell; bold dashed lines represent the example cell shown in (B). n denotes number of cells. For (D-G), asterisks indicate statistical significance (*p < 0.05; **p < 0.001).

To investigate the hidden mechanisms that enable the reversed directional preference in DSGC despite non-directional SAC response, we first tested if changes in SAC responses are reflected in DSGC inhibitory input. We voltage clamped DSGC to record their inhibitory (V_hold_ = 0 mV) and excitatory (V_hold_ = −60 mV) inputs in response to linear drifting gratings. In dark adapted conditions, DSGC inhibition was stronger during null than during preferred motion (Figure 4B, C). Comparison of the mean integrated inhibitory inputs during preferred and null motion over one grating cycle revealed ∼30% less total inhibition during preferred than during null motion (Figure 4D, top left). This asymmetric inhibition was also quantified by a negative amplitude DSI (DSI_DA_ = −0.22 ± 0.2; Figure 4E). Since linear motion in the null direction of a DSGC corresponds to CF motion in the SAC processes which innervate it, these findings matched SAC CF preference and were expected based on previous reports (Figure 2A-C) (Borst and Euler, 2011; Mauss et al., 2017; Vaney and Taylor, 2002; Wei and Feller, 2011). In accordance with SAC response kinetics (Figure 2D), kinetics of DSGC inhibition in response to preferred and null motion also differed, revealing a shorter rise time during null motion (Figure 4F). Examination of additional response parameters (fitted exponential power and response phase width) further supported consistency between SAC directional responses and DSGC inhibitory inputs (Supplementary Figure 4). Excitation to dark-adapted DSGC during preferred and null motion was on average equal in size (Figure 4D).

Following RVS, inhibitory inputs to DSGC dramatically differed in several ways compared to dark adapted conditions. First, inhibition was no longer stronger during null motion (Figure 4B, C). Comparison of the mean integrated inhibitory inputs during preferred and null motion demonstrated that overall inhibition in both directions became comparable (Figure 4D, bottom left). This loss of directional preference was also evident by the DSI values for inhibition, which clustered around zero (Figure 4E; DSI_RVS_ = 0.12 ± 0.2). Second, rise times during preferred and null motion turned comparable following RVS (Figure 4F). The described changes in inhibitory input to DSGC are in line with the observed loss of SAC CF preference following RVS (Figure 2, Supplementary Figure 4). Taken together, these results support the hypothesis that changes in inhibitory input to DSGC result from changes in SAC responses following RVS.

### Changes in timing of inhibition to DSGC underlie reversal of directional preference

As inhibitory inputs to DSGC became symmetric following RVS, we suspected that the RVS-mediated temporal shift in SAC responses underlies the reversal of directional preference in DSGC. To test this, we first assessed timing of synaptic currents in dark adapted conditions and following RVS based on the mean integrated inputs, comparing the time by which the evoked inputs reached half the integrated value (Figure 4D). We found that in dark adapted DSGC inhibition was faster during null than during preferred motion by ∼15 ms, while excitation was faster during preferred than during null motion by ∼35 ms. Following RVS, the converse was observed: inhibition was faster during original preferred than during original null motion by ∼40 ms, while excitation was faster during original null than during original preferred motion by ∼35 ms. Next, we measured the temporal differences between excitation and inhibition at the single cell level based on initial phases of synaptic inputs, set to the time the current waveform reached 20% of the response peak (see *Methods*). In dark adapted DSGC excitation tended to precede inhibition during preferred, but not null, motion (t_Exc_PD_-t_Inh_PD_ = −47 ± 66 ms; t_Exc_ND_ - t_Inh_ND_ = 36 ± 83 ms; p < 0.005), whereas following RVS inhibition tended to precede excitation during original preferred, but not original null, motion (t_Exc_PD_ - t_Inh_PD_ = 33 ± 77 ms; t_Exc_ND_ - t_Inh_ND_ = −43 ± 86 ms; p < 0.05; Figure 4B, G). The temporal shifts in excitation-inhibition balance support reduced response during original preferred motion and increased response during original null motion, in line with reversal of directional preference in DSGC spiking responses following RVS (Figure 4A). These findings match the observed surround-mediated phase shift in SAC directional response, indicating that the computation of motion direction in DSGC strongly depends on SAC center-surround receptive field organization which rules SAC response timing.

## Discussion

Here, we recruited RVS to dissect the role of center-surround organization in the computation of motion direction in the retina. SAC CF preference is thought to contribute to direction selectivity by providing DSGC with stronger inhibition during null motion. We found that RVS eliminates On-SAC CF preference and turns inhibition to DSGC symmetric. SAC CF preference as well as its loss following RVS could be predicted by a simple spatiotemporal model based on summation of the SAC responses to static rings, showing the dominance of SAC excitatory receptive field organization in shaping their directional response. Despite this loss of asymmetry in SAC directional responses following RVS, DSGC did not lose their ability to encode motion, but rather reversed their directional preference. The reversed computation originated from a shift in SAC response timing, which triggered a shift in the excitatory-inhibitory balance in DSGC that supported the null direction response. This time shift resulted from loss of SAC center and enhancement of its surround activation following RVS. Center-surround receptive fields can dynamically change with the visual input (Rivlin-Etzion et al., 2018; Wienbar and Schwartz, 2018), and such deviations in the receptive field organization of retinal ganglion cells were also detected *in vivo* (Sagdullaev and Mccall, 2005). Hence, SAC response timing can be adjusted in different contexts, imposing flexibility on the computation of motion in DSGC. This demonstrates a unique example of how anatomically-defined neuronal circuits can overrule their anatomy and dynamically change their function.

### SAC mediate direction selectivity via multiple mechanisms

SAC CF preference is a key mechanism in direction selectivity and is thought to provide larger inhibitory input to DSGC during null, but not preferred, motion. Different hypotheses were raised to account for SAC CF preference, including SAC intrinsic properties, inhibitory network connections and the specific distribution of their excitatory inputs (Borst and Euler, 2011; Demb, 2007; Mauss et al., 2017; Vaney et al., 2012; Wei, 2018; Wei and Feller, 2011). The input distribution hypothesis is supported by two recent studies showing that excitatory inputs are restricted to the proximal two-thirds of SAC processes and that this skewed distribution may enhance depolarization in the distal dendrites during CF motion (Ding et al., 2016; Vlasits et al., 2016). Moreover, it was previously suggested that spatiotemporal patterned excitation plays a role in SAC processes CF preference (Fransen and Borghuis, 2017; Greene et al., 2016; Kim et al., 2014). Our results support these findings, showing that dark-adapted SAC excitatory receptive field is constrained to its proximal dendrites and that a temporally-diverse excitation can predict SAC differential response to CF and CP motion. However, we suggest that this proximal excitation can serve DSGC direction selectivity not only by controlling the amount and kinetics of inhibition, but mainly by controlling its timing. During CF motion (corresponding to the DSGC original null direction) response quickly emerges in dark adapted SAC but arises in delay in SAC that were exposed to RVS. During CP motion (corresponding to original preferred direction) response is delayed in dark adapted SAC but arises earlier following RVS. Our experimental and modelling data suggest that this phase shift in SAC responses is mediated by SAC surround excitation, as SAC lose center responses following RVS. This demonstrates that SAC proximal excitation can support direction selectivity via multiple mechanisms, and emphasizes the supremacy of SAC response timing over its amplitude and shape in determining the directional output of DSGC.

Two recent studies aimed to abolish SAC CF preference to resolve its contribution to direction selectivity. Genetic elimination of SAC GABA-A receptors slightly reduced their CF preference and consequentially partially impaired DSGC directional tuning (Chen et al., 2016). Another study managed to impose symmetric inhibition to DSGC using the same genetic elimination of SAC GABA-A receptors, but only when photoreceptor inputs were pharmacologically blocked and SAC were activated using optogenetics (Hanson et al., 2019). Under these conditions, and despite the symmetric inhibition, direction selectivity to the original preferred direction is maintained in DSGC. In this case, the resilience of direction selectivity is the result of temporal asymmetries between inhibition and excitation provided by SAC, which corelease GABA and acetylcholine (Brecha et al., 1988; O’Malley and Masland, 1989; Vaney and Young, 1988). In our experiments, we utilized mechanisms that are naturally embedded in the network to impose symmetric inhibitory inputs to DSGC. Photoreceptors input was not blocked and excitation to DSGC emerged from a combination of glutamatergic and cholinergic inputs arising from bipolar cells and SAC, respectively. The change in SAC response timing following RVS probably shifted release time of both GABA and acetylcholine. Although SAC provide asymmetric inhibitory input to DSGC, SAC excitatory input to DSGC is symmetric – as SAC on both the preferred and null side form cholinergic connections onto the DSGC (Mauss et al., 2017; Vaney et al., 2012; Wei and Feller, 2011). As a result, the cholinergic input may cooperate with the GABAergic input to mediate null-direction computation following RVS: DSGC’s null motion corresponds with CF motion in null side SAC processes – resulting in delayed inhibition, but also corresponds with CP motion in preferred side SAC processes – resulting in faster excitation. Bipolar cells are also expected to shift their response timing since expansion of SAC receptive field results from enhanced surround activation in bipolar cells (Vlasits et al., 2014). Nevertheless, as bipolar cells and SAC bare different receptive field sizes, they probably differently change their response timing. Thus, temporal asymmetries between excitation and inhibition are not maintained following RVS, enabling the observed change in directional preference. Notably, our findings demonstrate that SAC response timing is not only necessary for the computation of direction, but is also sufficient to enable complete reversal in the computation of motion in the absence of other SAC response asymmetries.

### Center-surround antagonism and polarity switch in retinal cells

Like most retinal cells which bare opposite polarity preference to light in their center and surround, SAC display antagonistic center-surround organization (Fransen and Borghuis, 2017; Huang et al., 2019; Lee and Zhou, 2006; Peters and Masland, 1996). In dark adapted conditions, On-SAC hyperpolarized in response to a bright stimulus in the surround and depolarized in response to its disappearance (Figure 1D). Following RVS, On-SAC responses resembled these surround-mediated responses, even when stimulated in their center receptive field. This suggests that following RVS, On-SAC polarity switch originates from strengthening of their surround on the expense of their central response, in accordance with enhancement of antagonistic surround at high light levels (Barlow et al., 1957; Bisti et al., 1977; Dedek et al., 2008; Enroth-Cugell and Robson, 1966; Farrow et al., 2013; Rodieck and Stone, 1965). Polarity switch in On-SAC did not rely on changes in inhibitory surround circuits in the inner retina, but rather on opposed polarity activation of presynaptic On bipolar cells (Vlasits et al., 2014). This was suggested to result from loss of rod signaling, which takes part in the computation of motion direction (Rosa et al., 2016). Specifically, rods are saturated in response to the high-light levels RVS and no longer transduce light into electrical responses. Instead, they serve as relay neurons for cone-driven surround inhibition via horizontal cells (Rivlin-Etzion et al., 2018; Szikra et al., 2014; Vlasits et al., 2014). This surround activation is transferred from bipolar cells to SAC and can explain both the polarity switch and the expansion of SAC excitatory receptive fields from center to periphery. Other examples exist for antagonistic interplay between rods- and cones-mediated responses. Differential signaling from rods and cones was shown to generate antagonistic center-surround responses to mediate color opponency in mouse ventral retina (Joesch and Meister, 2016; Szatko et al., 2019). Here, too, horizontal cells signaling is thought to impose this antagonistic nature of rods and cones function via lateral inhibition.

### Null-tuned directional response and antagonistic surround

According to our data, SAC center and surround activation differentially affect their response timing, with temporal characteristics of SAC responses upon surround stimulation resemble temporal characteristics of SAC responses following RVS. Therefore, SAC center-surround organization is a crucial component in determining DSGC directional preference and acts upon it in a PD-center ND-surround antagonistic manner, supporting response to the preferred direction upon activation of its center but to the null direction upon activation of its surround. Antagonistic center-surround organization for motion encoding was previously described in the optic tectum of pigeons, where the cells’ response to motion in the center receptive field was facilitated by motion in the opposite direction in their surround (Frost and Nakayama, 1983). Similar effects were reported for direction selective cells in the cat superior colliculus and V1, as well as in area MT of primates (Allman et al., 1985; Von Grünau and Frost, 1983; Sterling and Wickelgren, 1969; Tanaka et al., 1986). We find a similar center-surround organization of opposing directions at the retina level. Such functional organization may act in the mouse direction selective circuit when exposed to high light levels that enhance surround activation. This hypothesis requires further investigation, but if true it implies that dynamic center-surround organization is beneficial for retinal encoding, not only for increasing sensitivity to fine spatial structures but also for sharpening any visual property encoded by retinal ganglion cells, including the computation of motion direction.

## Methods

### Animals

Two-photon targeted recordings from DSGC were performed using Drd4-EGFP (http://www.mmrrc.org/strains/231/0231.html) (Huberman et al., 2009) and Trhr-EGFP (http://www.mmrrc.org/strains/30036/030036.html) (Rivlin-Etzion et al., 2011) mice which express GFP in posterior-preferring On-Off DSGC. Recordings from On-SAC were performed using mGluR mice (Watanabe et al., 1998). Mice were from either sex, 4-7 weeks old. All procedures were approved by the Institutional Animal Care and Use Committee (IACUC) at the Weizmann Institute of Science.

### Electrophysiological Recordings

Mice were dark-adapted for at least 30 min before isoflurane anesthesia and decapitation. The retina was extracted and dissected in oxygenated Ames medium (Sigma, St. Louis, MO, USA) under dim red and infrared light. The isolated retina (dorsal part) was then mounted on a 0.22 mm membrane filter (Millipore) with a pre-cut window to allow light to reach the retina and put under a two-photon microscope (Bruker, Billerica, MA, USA) equipped with a Mai-Tai laser (Spectra-physics, Santa Clara, CA, USA) as previously described (Warwick et al., 2018). GFP cells were targeted for recordings with the laser set to 920 nm to minimally activate photoreceptors, and using a 60x water-immersion objective (Olympus, Tokyo, Japan). The isolated retina was perfused with warmed Ames solution (32–34 °C) and equilibrated with carbogen (95% O2:5% CO2).

Spike recordings from DSGC were made in loose cell-attached mode using 4–7 MΩ glass pipettes filled with Ames solution. Current-clamp recordings from SAC were made using 5–9-MΩ glass pipettes containing (in mM): 110 KCl, 2 NaOH, 2 MgCl2, 0.5 CaCl, 5 EGTA, 10 HEPES, 2 ATP, 0.5 GTP and 2 Ascorbate (pH = 7.2 with KOH; Osmolarity = 280). Voltage-clamp whole-cell recordings from SAC and DSGC were made using 5–9 MΩ glass electrodes filled with intracellular solution containing (in mM): 110 CsMeSO_3_, 2.8 NaCl, 4 EGTA, 20 HEPES, 5 TEA-Cl, 4 Mg-ATP, 0.3 Na3GTP, 10 Na2-Phosphocreatine and 0.025 AlexaFluor594 (pH = 7.2 with CsOH; Osmolarity = 290; E_Cl_ = −73mV). Holding voltages for measuring excitation and inhibition after correction for the liquid junction potential were 0 mV and −60 mV, respectively. Data were acquired using pCLAMP10, filtered at 2 kHz and digitized at 10 kHz with a MultiClamp 700B amplifier (Molecular Devices, CA, USA) and a Digidata 1550 digitizer (Molecular Devices).

### Light Stimuli

Visual stimuli were generated using Matlab and the Psychophysics Toolbox and were projected to the retina by a monochromatic organic light-emitting display (OLED-XL, 800 × 600 pixels, 85 Hz refresh rate, eMagin, Bellevue, WA, USA), through either 60x or 20x objectives (UMPLFLN60xW/UMPLFLN20xW; Olympus, Tokyo, Japan). All experiments were carried out in the photopic light range, with a background light intensity of 4.3×10^4^ R*/rod/sec (defined as light off) and a bright light intensity ranging between 4×10^5^ and 1×10^6^ R*/rod/sec. Visual stimuli were centered on the recorded cell’s soma and focused on the photoreceptor layer. For examination of SAC polarity, a bright spot (50 μm radius) on a dark background was presented to the cells for 2 sec. For investigation of SAC receptive field, 8 bright static rings (25 μm width) ranging between 25-200 μm radii were centered on cell soma and presented for 2 sec each, repeated five times in a pseudo-random order. SAC directional responses were assessed by presentation of expanding (centrifugal; CF) and collapsing (centripetal; CP) rings centered on the SAC soma, projected via the 20x objective for 5 sec (900 μm/sec, 2 Hz, 450 μm/cycle; first cycle was removed from analysis) and repeated five times in a pseudo-random order. SAC soma was masked by a 25 μm radius grey spot (Euler et al., 2002). DSGC directional responses were assessed from the responses to linear drifting gratings (900 μm/sec, 2 Hz, 450 μm/cycle; first cycle was removed from analysis) in 12 different directions, projected via the 60x objective for 3 sec and repeated four times in a pseudo-random order. Repetitive visual stimulation (RVS) consisted of linear gratings (900 μm/sec, 2 Hz, 450 μm/cycle) moving in the preferred and null directions for 40 sec and repeated up to 4 times.

### Computational Modeling

We reconstructed an individual SAC filled with AlexaFluor594 using Neurolucida 360 Studio from a three-dimensional volume data acquired using a two-photon microscope (Bruker, Billerica, MA, USA) at 760 nm. Images were acquired at 0.5 μm interval using a 60x objective (UMPLFLN60xW Olympus, Tokyo, Japan). We assumed SAC processes were completely flat and therefore used a projection of the reconstructed SAC. We estimated the diameter of each dendritic region based on its distance from the soma according to values acquired from EM data: 0.2*e ^-x/40^ + 0.15 (Poleg-Polsky et al., 2018).

The SAC was discretized to 755 segments. To mimic the different excitatory receptive fields, we implanted excitatory inputs in the form of synapses in a semi-random pattern according to a predefined uniform density distribution. The density distribution was set to 12% of the sections at the proximal 2/3 or distal 2/3 of the dendritic tree, generating 238 and 528 synapses for dark adapted conditions and following RVS, respectively. Ribbon synapses were modelled using current clamps, each bound to a postsynaptic channel defined by a ligand-activated Markov sequential-state machine, where the ready releasable pool of vesicles was set to 70, generating a maximal post synaptic current of 200 pA, with a 5% probability of release and refilling rate of 4 vesicles per simulation interval (25 ms) (Singer and Diamond, 2006). Membrane ion channels were defined as: Rm = 10 kOhm/cm^2^; Cm = 1 μF; Ri = 75 Ohm; Vm = −60 mV; Channels density (in S/cm^2^): NaV1.8 0.25e^-3^; Kdr at the soma 3e^-3^, 2e^-3^ at the processes; L-type Ca^2+^ 1e^-3^ at the distal 2/3 and zero elsewhere (Ding et al., 2016).

Visual stimuli were introduced to the simulated SAC via the excitatory synapses. This was accomplished by current-clamping a presynaptic compartment that represented each synaptic input according to the spatiotemporal pattern of the stimuli. Visual stimulation resembled the CF and CP moving rings (900 μm/s, 2 Hz, 450 μm/cycle). Simulations were conducted using NEURON 7.6 and run on Intel i9-7900x 3.3GHz 10-core Processor, 64GB RAM and NVIDIA Titan Xp 12GB GPU, with a Linux operating system. Simulated SAC responses were taken from SAC soma. For dendritic voltage measurements we randomly picked a distal dendrite (d=135 μm from soma).

### Quantification and statistical analysis

We extracted spike times from the data after offline filtration using a 4 pole Butterworth bandpass filter between 80 and 2000 Hz. The preferred direction of each cell in dark adapted conditions was determined by first normalizing the average spike count in each direction by the total number of spikes for all directions. The vectorial summation of these normalized responses yielded a vector which direction was the preferred direction of the cell and which magnitude gave the width of the tuning (vector sum, ranges between 0 and 1). The direction selective index (DSI) for each cell was calculated as:

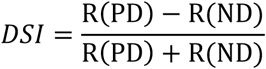

where *R(PD)* is the average number of spikes in the stimulus direction closest to the preferred direction and *R(ND)* is the average number of spikes in the stimulus 180 degrees opposite. In dark adapted conditions DSI values were positive, reaching a maximum of 1 for perfectly tuned DSGC. Following RVS the DSI was calculated based on the original preferred and null directions, resulting in negative values for DSGC which reversed their directional preference. In cells that were recorded only following RVS (i.e., were exposed to RVS before recording started), the preferred direction was set according to at least one dark adapted cell within the same retinal piece which determined the posterior direction of the preparation (assuming all GFP positive cells were tuned to the same posterior direction) (Huberman et al., 2009; Rivlin-Etzion et al., 2011).

Voltage-clamp raw traces were processed with a Savitzky-Golay filter (order: 3; frame length: 81). Excitatory current traces were inverted in sign. To assess DSGC response phases, baseline activity was determined based on minimum value of mean trace. Phases with an amplitude greater than baseline activity + 2 STD were detected based on local maxima. The phase duration was set according to the number of bins above threshold surrounding the local maxima and phases shorter than 50 ms were excluded from analysis. Phase onset was determined as the time the response reached 20% of the response peak and phase rise time was determined as the time from phase onset to phase response peak. For determining the time difference between excitatory and inhibitory phases, we first identified the max phase in each holding potential (based on amplitude) and then matched phases which were separated by less than 250 ms. The time difference between matching excitatory and inhibitory phases was measured as the difference between their response onsets. Directional tunings of inhibitory currents were determined by the DSI, using the current amplitudes during PD and ND motion.

SAC directional tuning was determined by:

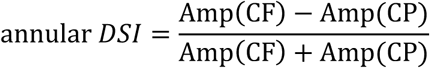

where *Amp(CF)* and *Amp(CP)* are the amplitudes of the SAC’s mean response to CF and CP motion. To quantify response delay, we assessed the time from stimulus onset to response onset (time the response reached 20% of the response peak). In dark adapted conditions, we determined the stimulus onset as the time the leading bright edge of the ring encountered the cell’s dendritic arbor (i.e., when leading edge traversed most proximal dendrites for CF motion and most distal dendrites for CP motion). Following RVS, we used similar measures but in relation to the dark edge of the ring, due to SAC polarity switch. For CF motion, we took into consideration the masking of SAC soma with a 25 μm radius spot (Euler et al., 2002) for determining stimulus onset. Response onset and rise time were calculated as described above for DSGC’s synaptic inputs.

Simulation of SAC responses to motion is based on their responses to static rings of different radii, where each mean response was shifted in time to simulate motion-induced activation (Lien and Scanziani, 2018). In the dark-adapted condition we set time 0 to 200 ms prior to ring onset, following RVS it was set to 200 ms before ring offset due to polarity switch. To mimic CF motion, responses started with the most proximal ring and ended with the most distal ring, with each consecutive ring response being shifted by additional 27 ms (to mimic ∼900 μm/sec speed). For CP motion, responses started with the most distal ring and ended with the most proximal ring. The shifted waveforms were then summed to predict the motion response.

For statistical analysis, data sets were tested for normality using Chi-squared goodness of fit test. Data sets that followed normal distribution were compared using a two-sample unpaired or paired Student’s t-test, according to data structure and Wilcoxon rank sum test was used for abnormally distributed data sets. Throughout the figures, sample statistics and average waveforms are expressed as means ± SEM, unless specified otherwise.

## Supplementary Figures

**Supplementary Figure 1:**
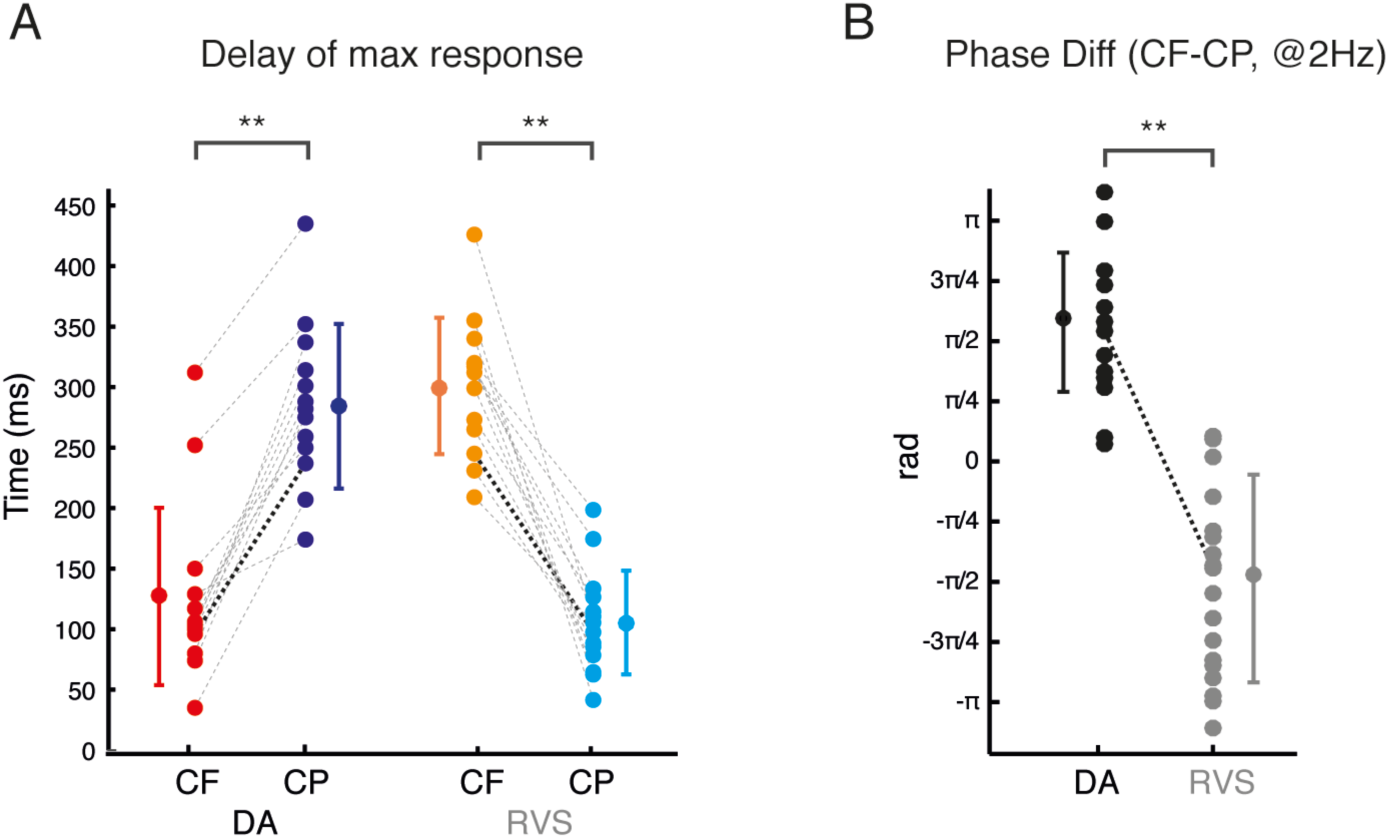
RVS-mediated phase shift in On-SAC response is evident in different quantification methods. (A) The delay to response peak in On-SAC, measured as the latency from the time the bright and dark rings appeared on the cell’s receptive field to response peak time in dark adapted conditions and following RVS, respectively. (B) Phase differences between responses during CF and CP motion in dark adapted conditions (DA, black) and following RVS (RVS, grey), calculated based on the difference of the Fourier transform argument at 2 Hz frequency. Asterisks indicate statistical significance (**p < 0.001).

**Supplementary Figure 2:**
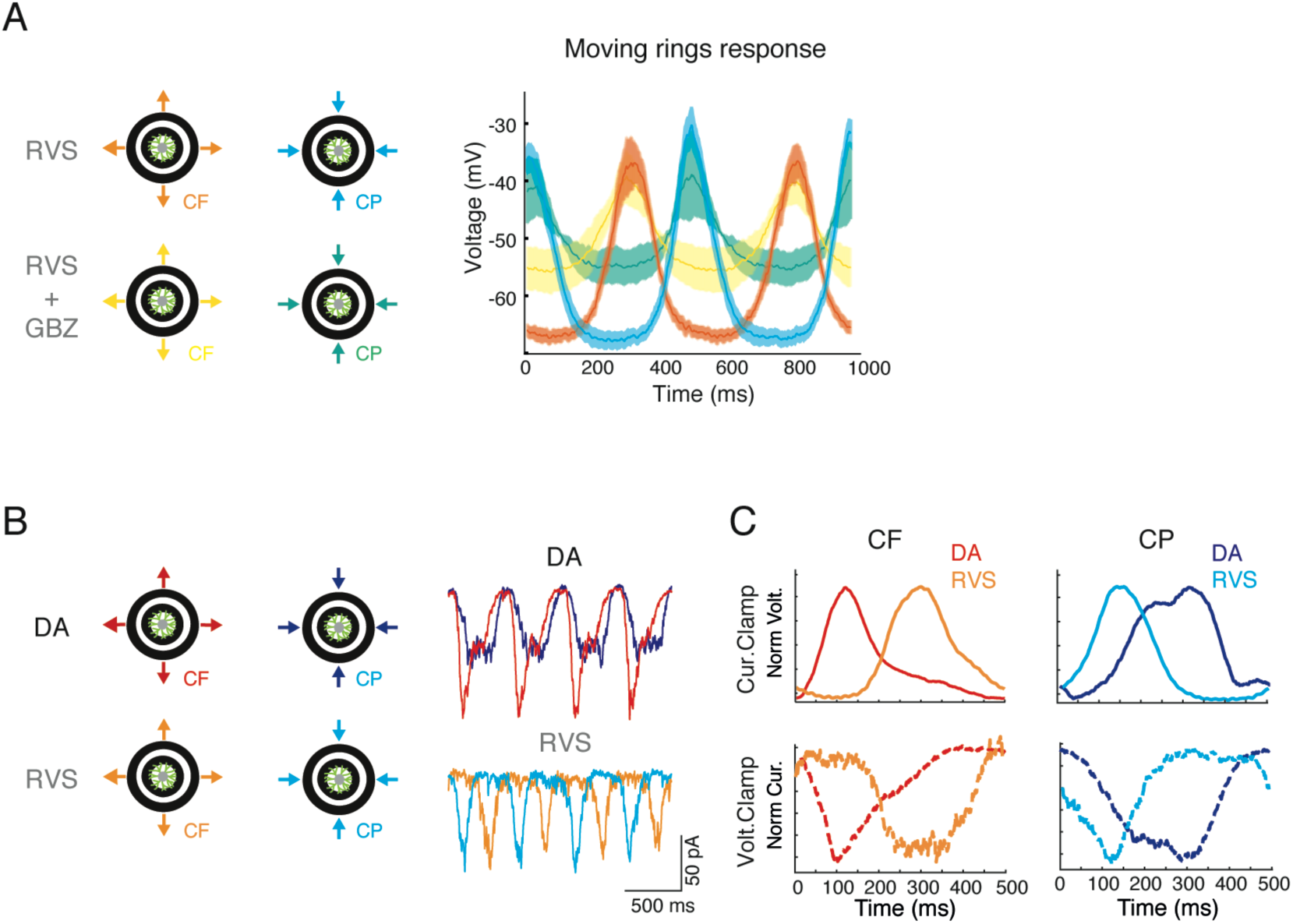
On-SAC response time shift is independent of GABAergic inhibition and is reflected in their excitatory currents. (A) *Left:* Illustration of CF and CP motion presented to On-SAC following RVS and after application of gabazine (10 μM). *Right:* Gabazine application does not change the phase of the response but abolishes the hyperpolarizing response to bright ring motion. (B) *Left:* Illustration of CF and CP motion presented to On-SAC in dark adapted conditions and following RVS. *Right:* Average voltage clamp recordings of the excitatory currents (V_hold_ = −60 mV) of an example On-SAC in response to CF and CP motion in dark adapted conditions (DA, top) and following RVS (RVS, bottom). (C) On-SAC population normalized voltage (top) and excitatory current (bottom) waveform average in response to CF (left) and CP (right) motion. The responses are shifted according to the cell’s polarity preference, with time 0 denoting the appearance of the bright and dark ring in dark adapted conditions and following RVS, respectively. Note that On-SAC depolarization measured in current clamp mode resembles in time excitatory currents measured in voltage clamp mode, indicating that time shifts in On-SAC following RVS are mediated by changes in excitation.

**Supplementary Figure 3.**
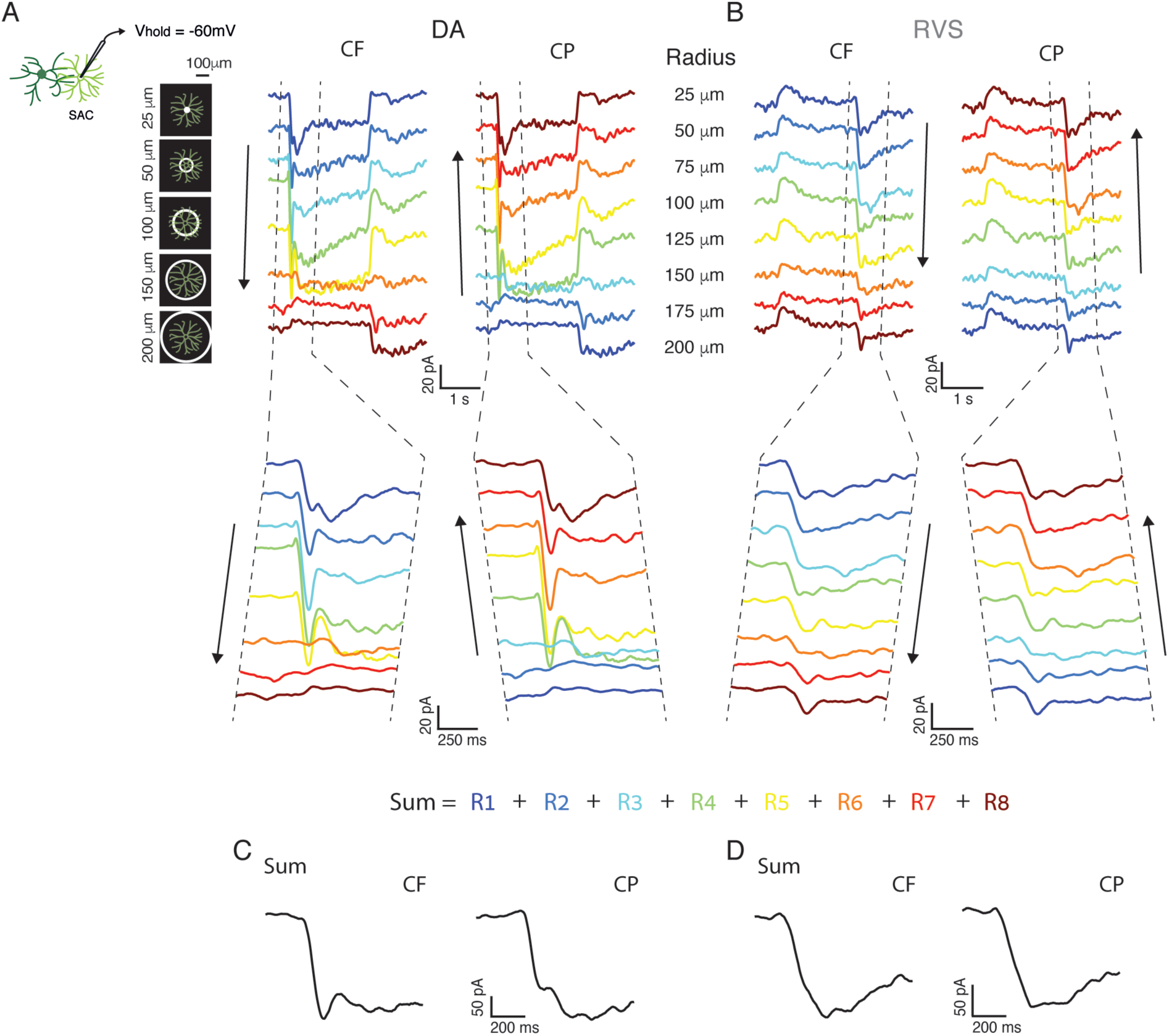
Direction selectivity in On-SAC can be explained by linear summation of their responses to static stimuli. (A) Population average of EPSCs (Vhold = −60 mV) evoked in dark adapted SACs in response to rings of different radii (several rings are illustrated on the left; all rings radii are marked on the right). The traces are color coded by their time shift, where the blue trace is not shifted at all and each consecutive trace in shifted in time by 27 ms from its previous (with brown trace shifted by 7×27 ms = 189 ms). CF motion simulation is depicted on left and CP motion simulation is depicted on right. Tilted vertical lines indicate the time window used for linear summation of the responses. *Bottom*: magnification of the time window used for linear summation in (C) and in Figure 3D. (B) As in (A), but for SAC recorded following RVS. *Bottom*: magnification of the time window used for linear summation in (D) and in Figure 3E. (C) The resulting linear summation of the responses to CF-like motion (left) and to CP-like motion (right). (D) As in (C), but for SAC recorded following RVS.

**Supplementary Figure 4.**
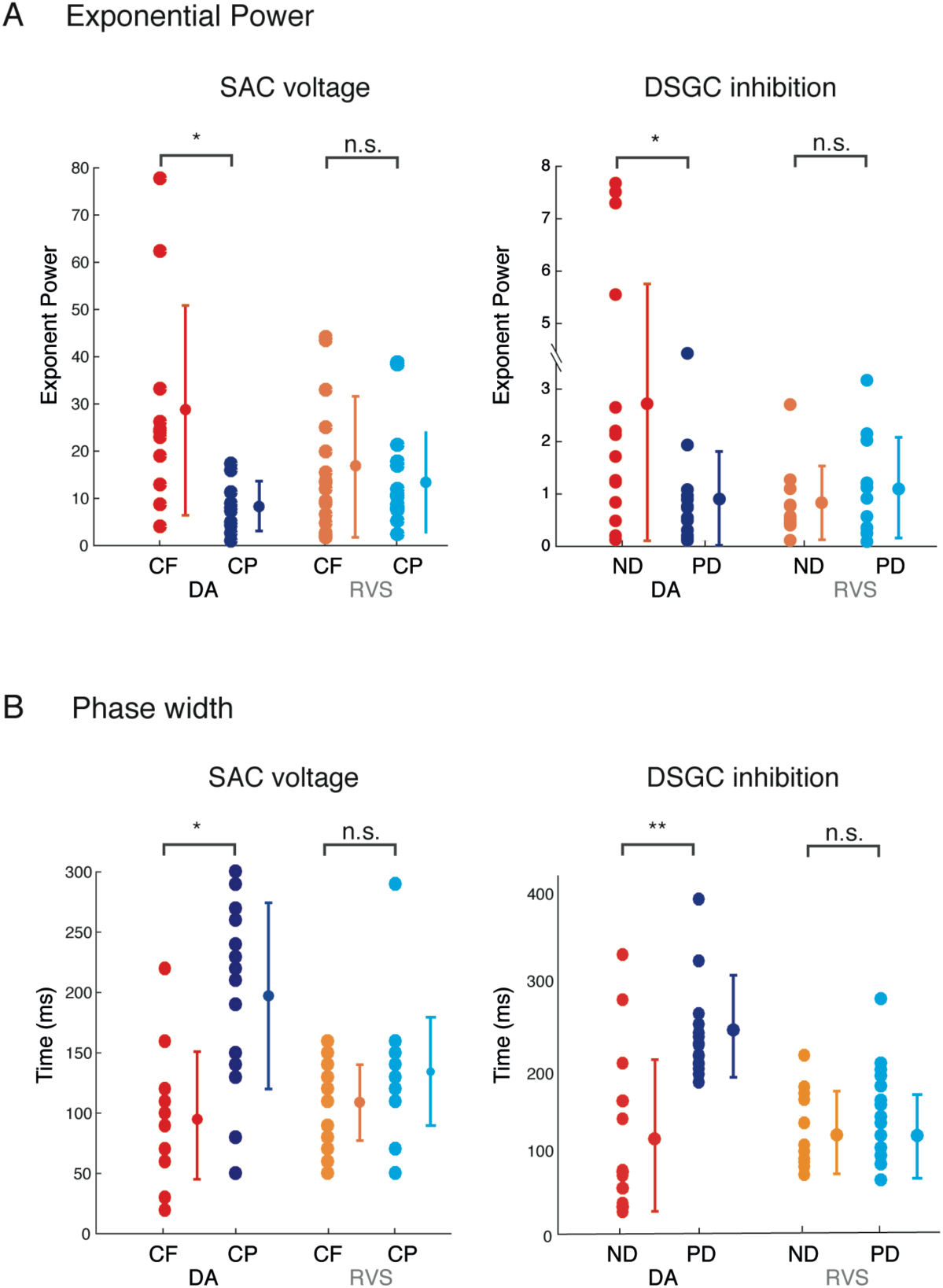
On-SAC response and DSGC inhibition are comparable across quantification methods. (A & B) Comparison between SAC voltage responses to CF and CP motion (left) and DSGC inhibitory currents to linear drifting gratings in the null (ND) and preferred (PD) directions. (A) depicts the exponential power of the best fitting curve (highest r^2^) to the rising of the responses. (B) Depicts response width, calculated as the difference between the maximum (positive slope) and minimum (negative slope) points in the waveform’s derivative. Asterisks indicate statistical significance (*p < 0.05; **p < 0.001). Note the similarities between On-SAC CF and DSGC ND parameters, as well as On-SAC CP and DSGC PD parameters in both dark adapted conditions and following RVS. Compare also main Figure 2C with main Figure 4E and main Figure 2D with main Figure 4F.

## Acknowledgements

We thank Marla Feller, Anna Vlasits, E.J. Chichilnisky, Ilan Lampl, Yaniv Ziv, Michal Schwartz and members of the Rivlin lab for useful comments and discussions. The authors acknowledge support from the I-CORE (51/11), the Minerva Foundation, the ISF Foundation (1396/15), the European Research Council (ERC-StG 757732), Dr. and Mrs. Alan Leshner, the Lubin-Schupf Fund for Women in Science, the Charles and David Wolfson Charitable Trust, and Ms. Lois Pope. L.A is supported by the Israeli Education Foundation (ISEF). M.R.-E. is incumbent of the Sara Lee Schupf Family Chair.

